# gEAR: gene Expression Analysis Resource portal for community-driven, multi-omic data exploration

**DOI:** 10.1101/2020.08.28.272039

**Authors:** Joshua Orvis, Brian Gottfried, Jayaram Kancherla, Ricky S. Adkins, Yang Song, Amiel A. Dror, Dustin Olley, Kevin Rose, Elena Chrysostomou, Michael C. Kelly, Beatrice Milon, Maggie S. Matern, Hela Azaiez, Brian Herb, Carlo Colantuoni, Robert L. Carter, Seth A. Ament, Matthew W. Kelley, Owen White, Hector Corrada Bravo, Anup Mahurkar, Ronna Hertzano

## Abstract

The gEAR portal (gene Expression Analysis Resource, umgear.org) is an open access community-driven tool for multi-omic and multi-species data visualization, analysis and sharing. The gEAR supports visualization of multiple RNA-seq data types (bulk, sorted, single cell/nucleus) and epigenomics data, from multiple species, time points and tissues in a single-page, user-friendly browsable format. An integrated scRNA-seq workbench provides access to raw data of scRNA-seq datasets for *de novo* analysis, as well as marker-gene and cluster comparisons of pre-assigned clusters. Users can upload, view, analyze and privately share their own data in the context of previously published datasets. Short, permanent URLs can be generated for dissemination of individual or collections of datasets in published manuscripts. While the gEAR is currently curated for auditory research with over 90 high-value datasets organized in thematic profiles, the gEAR also supports the BRAIN initiative (via nemoanalytics.org) and is easily adaptable for other research domains.

## INTRODUCTION

Multi-omic genome-wide functional profiling of gene expression, epigenomic modifications, and genome accessibility have evolved as the workhorse for discovery in biological sciences. Gene expression data range from bulk to single cell/nucleus RNA-seq (scRNA-seq/snRNA-seq, respectively), ATAC-seq for genome accessibility, and methyl-seq, ChIP-seq and CUT&RUN to profile the epigenome^1^. Furthermore, samples range in number from few to thousands of bulk RNA-seq samples (e.g., GTEx^2^) or, in the case of single cell assays, may include hundreds to millions of individual cells. These data, generated by individual laboratories or consortia, are used both as hypothesis-generating tools and in comprehensive atlases where they are intended for re-use^3–5^. While often deposited in public repositories such as the Gene Expression Omnibus (GEO)^6^, sequence read archive (SRA)^7^, or even domain specific repositories such as Synapse (synapse.org) – this kind of access requires download of the data, and often re-analysis and curation, which are time-consuming and frequently outside the skillset available to many biologists. The full potential of these resources is not realized unless meaningful access is available to all potential users. Thus, an even greater challenge than the unending struggle to handle the increasing scale of data produced by omics technologies is that of creating, curating and populating responsive, scalable visual analysis tools with the data they produce.

A number of web-based viewers for gene expression data are now available, including the Single Cell Expression Atlas^8^, ShinyOmics^9^ or mixOmics^10^, cBioPortal (https://www.cbioportal.org/), Xena (https://xena.ucsc.edu/) and the Single Cell Portal (SCP) (https://singlecell.broadinstitute.org/single_cell). However, current web-based viewers have limitations that have prevented their broad adoption. For example, the SCP enables users to visualize the expression of genes in the context of cell clusters and cell types from a small but growing number of scRNA-seq datasets. The number of display types is limited, interactive analysis is reserved to precomputed steps already performed on raw data, and data types other than scRNA-seq are not supported. Other tools such as ShinyOmics^9^, CellxGene (cellxgene.cziscience.com) or mixOmics^10^, for example, are part of a growing number of applications that aim to support end-users who perform analyses of their own datasets but are designed to analyze one dataset at a time^11^. Some tools assimilate data from several studies for cross-normalization and further analysis, but individual source datasets lose their original context in these cases (https://www.cancer.gov/tcga). Notably, few if any tools allow the display of results from more than one study at a time, which is desirable for comparison.

The gEAR (gene Expression Analysis Resource) is an open access online platform that integrates web-based visualization and analysis abilities for multi-omic data and a user-friendly single-cell RNA-seq analysis workbench. The gEAR is designed to serve as a community driven research resource for biologists from different backgrounds interested in functional genomics data. In the implementation presented in this manuscript, the gEAR aims to make multi-omic data easily accessible for the hearing research community to facilitate data mining, enable comparative multi-omic analyses, and encourage data sharing thereby helping advance auditory research for future therapeutic discoveries for hearing loss. The gEAR currently contains over a trillion hearing and balance related datapoints from more than 400 gene expression datasets generated by different platforms (RNA-seq, scRNA-seq, epigenetic and microarray datasets), of which 170 are public. However, data from any domain can be uploaded to the gEAR and clones can be developed and customized to support other research communities. gEAR has already been deployed to support the BRAIN initiative (nemoanalytics.org) and the Institute for Genome Sciences (IGS) Genomic Centers for Infectious Diseases (gcid.umgear.org).

## RESULTS

### The gEAR portal overview

The gEAR directly advances discovery in the fields of hearing and balance research, supports collaboration and empowers biologists to better perform their work. It is designed to serve biologists interested in functional genomics data relevant to their research and requires little previous knowledge of bioinformatics tools and no programming expertise. The gEAR consists of six main components: (1) A dataset uploader and curator allow users to upload their own data and choose from a variety of displays to visualize their data including interactive bar, line, scatter or violin plots; colorized anatomical graphical representations of the data; and ordination plots such as tSNE and PCA plots; (2) A dataset manager used to group and arrange datasets in profiles, and to specify which individual or groups of datasets are shared; (3) A gene expression browser to visualize gene expression that also provides value-added resources such as functional annotation and link-outs to relevant repositories; (4) A basic comparison tool to compare and visualize expression between any two conditions of the same dataset (e.g., phenotypic groups, timepoints); (5) A single-cell workbench to perform *de novo* analyses from scRNA-seq count data or start with existing expertly analyzed data, and perform additional analyses and visualizations; and (6) Integration with Epiviz^12^, a linear genomic browser which allows simultaneous visualization of genome accessibility and epigenetic information alongside the expression results; (**Fig. 1**).

**Fig. 1:**
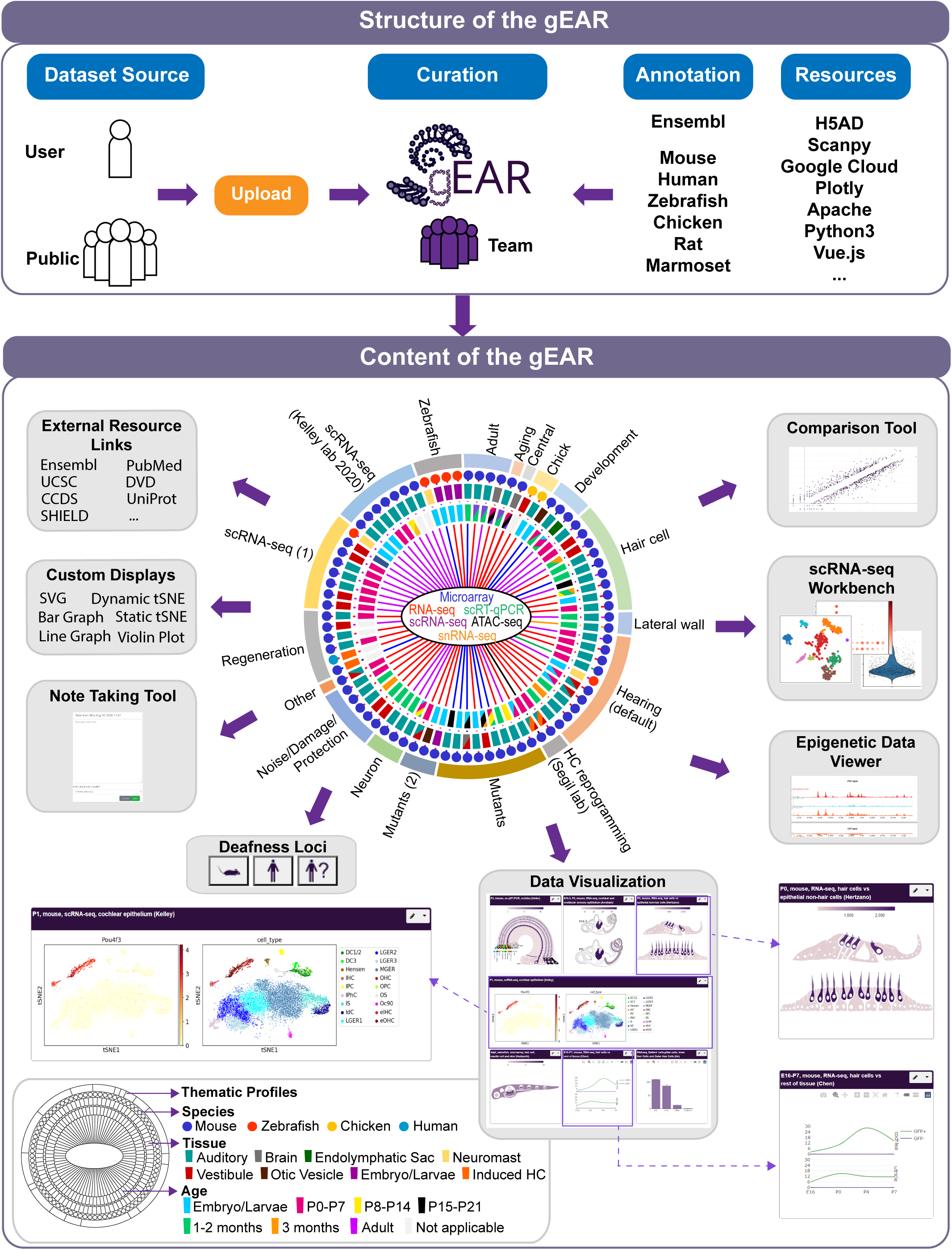
gEAR overview. The gEAR is a community-based portal for visualization, analysis and sharing of multi-omic data. Data consist of contributions from individual users as well as public data uploaded and curated by the gEAR team. Users can view multiple datasets from a variety of organisms, methods, time points or tissues in the same page. Each dataset is linked to informatic tools for additional annotation, data exploration (e.g., a compare tool, and a single cell analysis workbench) and an ability to customize the display based on available metadata. Datasets link also to the GEO entry, publication and a note taking tool. The gEAR is current curated for the ear field where over 400+ datasets from 6 species and a variety of methodologies are displayed in thematically curated profiles (collections of datasets).

### gEAR viewer: simultaneous exploration of multiple datasets and data modalities

One of the functionalities of the gEAR is the simultaneous exploration and visualization of gene expression in multiple datasets, which represent data from different modalities or organisms (**Fig. 1**). Searching for a gene on the gEAR main page yields a grid of results displayed as cells/boxes each corresponding to the gene’s expression in a single dataset. We call the selection and arrangement of these datasets a ‘profile’, and the gEAR currently has over 90 such datasets curated into 18 thematically related profiles (e.g., hearing, regeneration, damage, etc.). Furthermore, registered users can create their own named profiles and populate them with public as well as their own private datasets. Creating multiple topic-specific dataset profiles can be a powerful way for a user to interrogate expression data across conditions (e.g., is a gene that changes in response to noise also differentially expressed in a published mutant line) and validate their own experimental results by comparing them to published data before publication.

#### The gEAR system supports rapid and meaningful data exploration

Dataset banners succinctly describe the data and contain a drop-down menu to allow for easy access to many of gEAR’s built-in analysis tools and links to additional external information, including the reference publication. Users can also record notes on datasets, marking the notes private or public. The gEAR main results page offers users added value such as expandable organism-specific gene annotation, and useful external resource links which can be general (e.g., to PubMed and UCSC^13^) or domain specific (e.g., DVD - Deafness Variation Database^14^). In addition, domain-specific features can be added as a plug-in, such as Deafness gene annotation in the case of the gEAR adaptation for the hearing research community. Here if a gene is a known deafness causing gene in human or in mouse, or maps to a deafness locus that has yet to be cloned, a corresponding highlighted icon provides users further links to the appropriate resource (e.g., OMIM^15^, MGI (http://www.informatics.jax.org/), IMPC^16^ or a manuscript describing the locus) (**Fig. 1**).

*The Dataset Manager tool* (**Supplementary Fig. 1**) allows users to explore which datasets are available in the system, select datasets to build a profile, and customize their display. With the Dataset Manager, users can also generate short private URLs to share individual datasets or entire profiles with collaborators while keeping them otherwise hidden from public view. When researchers publish their own work, users are provided with a permanent URL (Permalink), that will be maintained over time to allow them to link the data to the manuscript and disseminate the data in a meaningful and browsable way (**Supplementary Fig. 1**). A common user workflow consists of 1) uploading the data privately when its initially received, 2) using the gEAR to interrogate the private dataset and view it alongside other published ones and 3) when the data are ready to be published, including a permalink URL in the paper and toggling the dataset’s access to public upon acceptance.

### gEAR curator: Meaningful data exploration through meta-data based customizable displays

The best insights into biological data can be gleaned when users can tailor visualizations to their analysis needs as much as possible. Support for visualization in the first versions of gEAR was to colorize scalable vector graphics (SVG) of anatomical sites rendered as cartoons, where each cell or structure is colored based on a user-customizable color gradient derived from the dataset expression matrix (**Supplementary Fig. 2**). For example, single-cell expression data mapped onto the cellular anatomy of the cochlea (the inner ear hearing organ) allow researchers to appreciate the gradual attenuation of gene expression in cells from the apex to base of this organ (**Supplementary Fig. 3**). Hovering over an anatomical site displays expression levels with a tooltip while a gradient bar at the top of the dataset displays similar information.

The gEAR framework has since been expanded to support interactive bar charts, time series, violin plots, scatter plots, ordination plots such as tSNE, UMAP and principal components, and multi-graph capabilities (**Fig. 2**). The properties of each dataset define the offered displays. gEAR displays support grouping data based on experimental metadata (e.g., data separated by sex, time point or aggregated, **Fig. 2a**). For scRNA-seq the gEAR offers two additional specialized types of display. Dynamic scatter plots (e.g., tSNE/UMAP) allow users to customize the coloring of the display and its tool-tip (hover-over) annotation based on available metadata; static scatter plots are also available to support faster rendering of very large datasets and side-by-side visualization of ordination plots colored by expression and cluster or other selected metadata (**Fig. 2b**). Importantly, many datasets, in particular scRNA-seq, lend themselves to SVG-based representations for intuitive visualization of data, resulting in impressionist-style images (**Fig. 2c**). Multiple easily interchangeable displays can be generated for any datasets using the Dataset Curator, offering additional insight into biological phenomena (**Supplementary Fig. 4**). Furthermore, when logged into the portal, data are displayed according to the users’ preferences allowing further flexibility and adaptability to individual user’s needs.

**Fig. 2:**
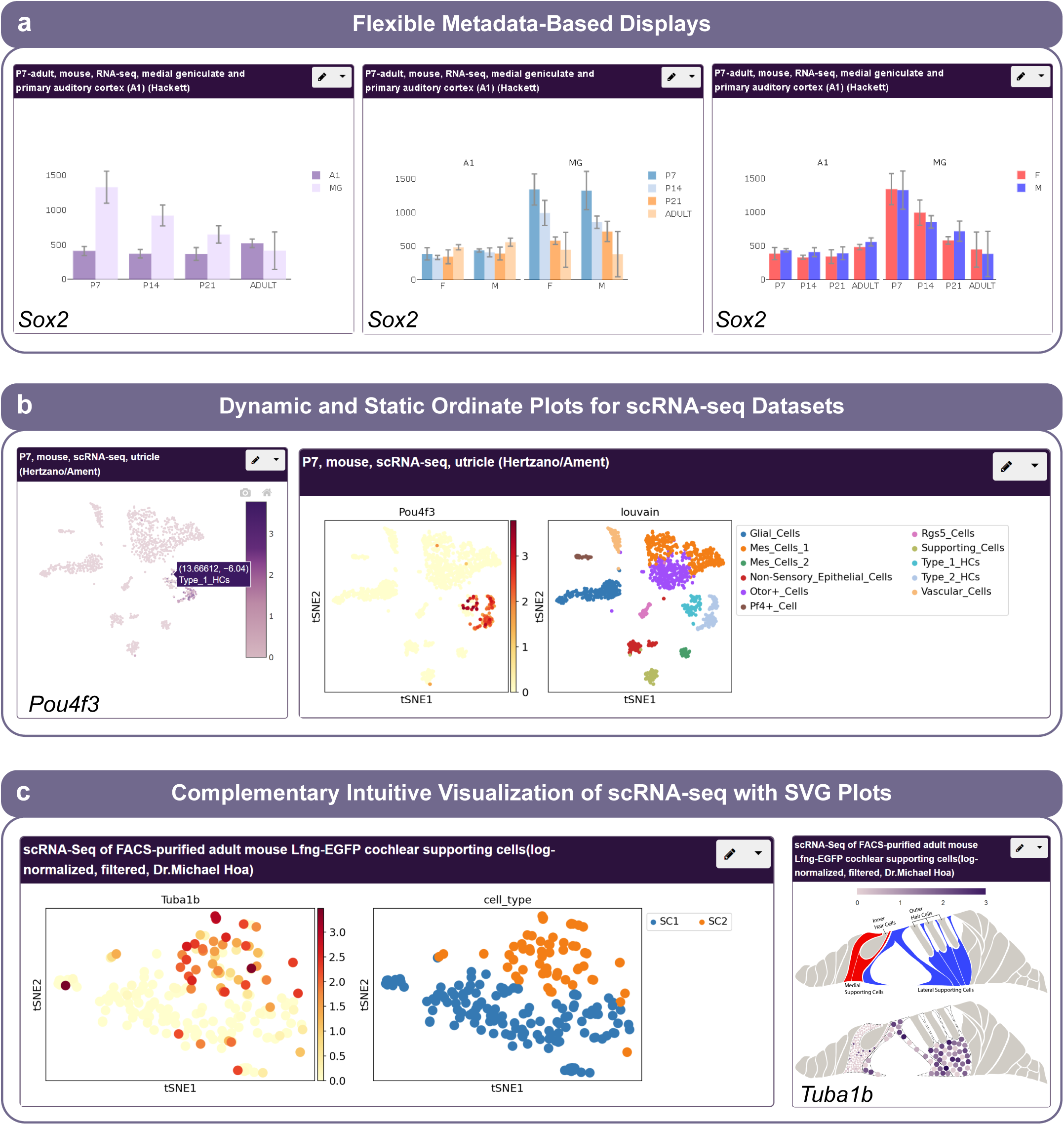
gEAR Data Visualization Options. **a**, Flexible metadata-based graph displays (here, parsing the data by cell type and/or sex and time point^65^). Left panel: data sorted by time point and tissue. Middle and right panels: graphs separated also by sex demonstrating two displays highlighting different aspects of the data. **b**, Data displays for scRNA-seq. Left panel: dynamic tSNE – hovering over any cell indicates cell type and expression level. Right panel: static tSNE with a side-by-side plot representing user-selected metadata. **c**, scRNA-seq data presented both in an ordinate plot and in an SVG-based visualization, here demonstrating the added value of this form of data display by improving ease of cell source identification^24^.

### gEAR Data Analysis: tools for bulk and single cell data exploration

The power of publicly available omic data is in its ability to be used as a resource. New knowledge is generated by custom analyses prompted by user questions. Thus, to empower users to meaningfully interrogate data, tools for rapid identification of differentially expressed genes, marker genes and other comparisons are necessary.

#### The gEAR compare tool

The simplest use case for data exploration is differential expression between conditions/cell types within a dataset. The compare tool allows two dataset conditions to be compared via an X-Y scatter plot (**Fig. 3**). This is appropriate for bulk RNA-seq or scRNA-seq datasets which have pre-computed clusters, conditions or cell type assignments. The scatter plot itself is interactive, displaying additional information such as the expression value and gene symbol of hovered-over data points. Users can control which genes are displayed using fold-change cutoff and standard deviation filters (**Fig. 3a**). If the data are uploaded with experimental design annotation indicating multiple replicates per sample, statistical significance is determined by using a t-test or Wilcoxon-Rank-Sum. Differentially expressed genes can be selected and downloaded as plain text files, or saved as gene carts, to be used anywhere else within the gEAR where a group of genes can be used as input (**Fig. 3b**).

**Fig. 3:**
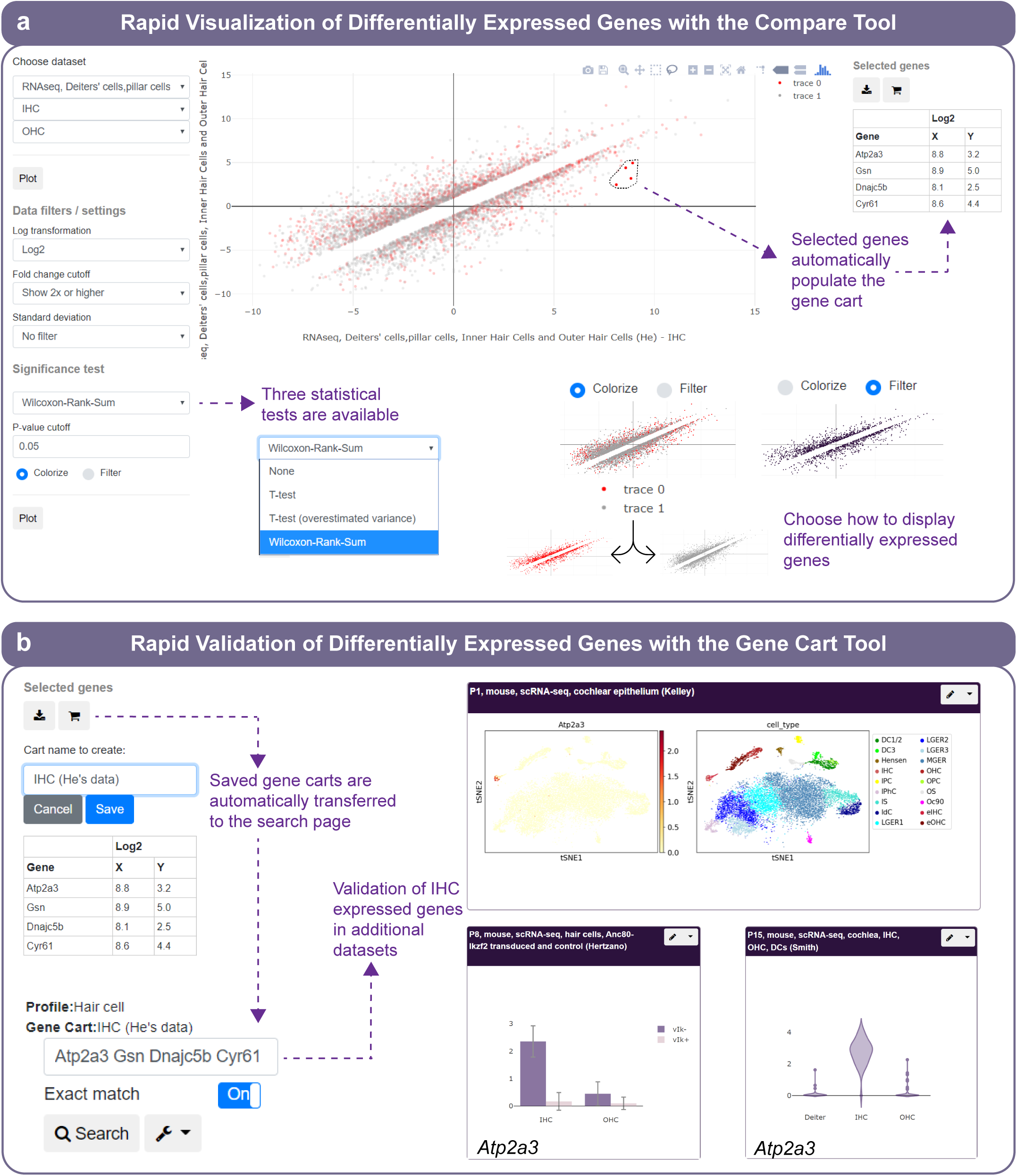
The gEAR compare tool. **a**, Result page for the comparison of inner hair cells (x-axis) versus outer hair cells (y-axis) of a public dataset^53^. Users can elect to apply a statistical test to the scatter plot comparing data across conditions. Genes can then be colorized based on significance, or non-significant genes can be filtered out. Users can select genes of interest (here, highly expressed in IHC) to appear in a downloadable table. **b**, The list of selected genes is saved as a cart and used as input to search other datasets. The high expression in IHC of the selected genes is confirmed in three independent datasets^5,23,66^. This figure is an example of the added value for immediate validation of data exploration by having a centralized resource curated data from multiple related sources.

#### The gEAR scRNA-seq Workbench

A popular and feature-rich tool of the gEAR is the single-cell workbench which allows users to perform *de novo* analysis of an existing or new dataset (**Fig. 4a**) or explore existing pre-loaded analyses (e.g., an uploaded Seurat object^17^ or any dataset with pre-defined clusters/cell types, **Fig. 4b**). When performing a *de novo* analysis, the workbench visually guides the user through a full, custom analysis. The single cell workbench is largely based on the Scanpy^18^ pipeline implemented in a graphical user interface (GUI) offering enhanced visualization options and faster data processing than standard R-based pipelines due to the use of dynamically-scalable Google cloud instances and data storage in binary H5AD-formatted files (http://www.hdfgroup.org/HDF5/). An analysis includes initial quality control steps, differential expression analysis, principal component analysis, dimensionality reduction (tSNE and UMAP), clustering, marker gene identification, and cluster comparison visualization. Users can access this tool both from the front page and any dataset panel by choosing the “Analyze” link. Analyses generated by users are automatically stored and can be named for later retrieval. User-generated analyses can also be made public or used to curate datasets for visualization in the gene browsing page.

**Fig. 4:**
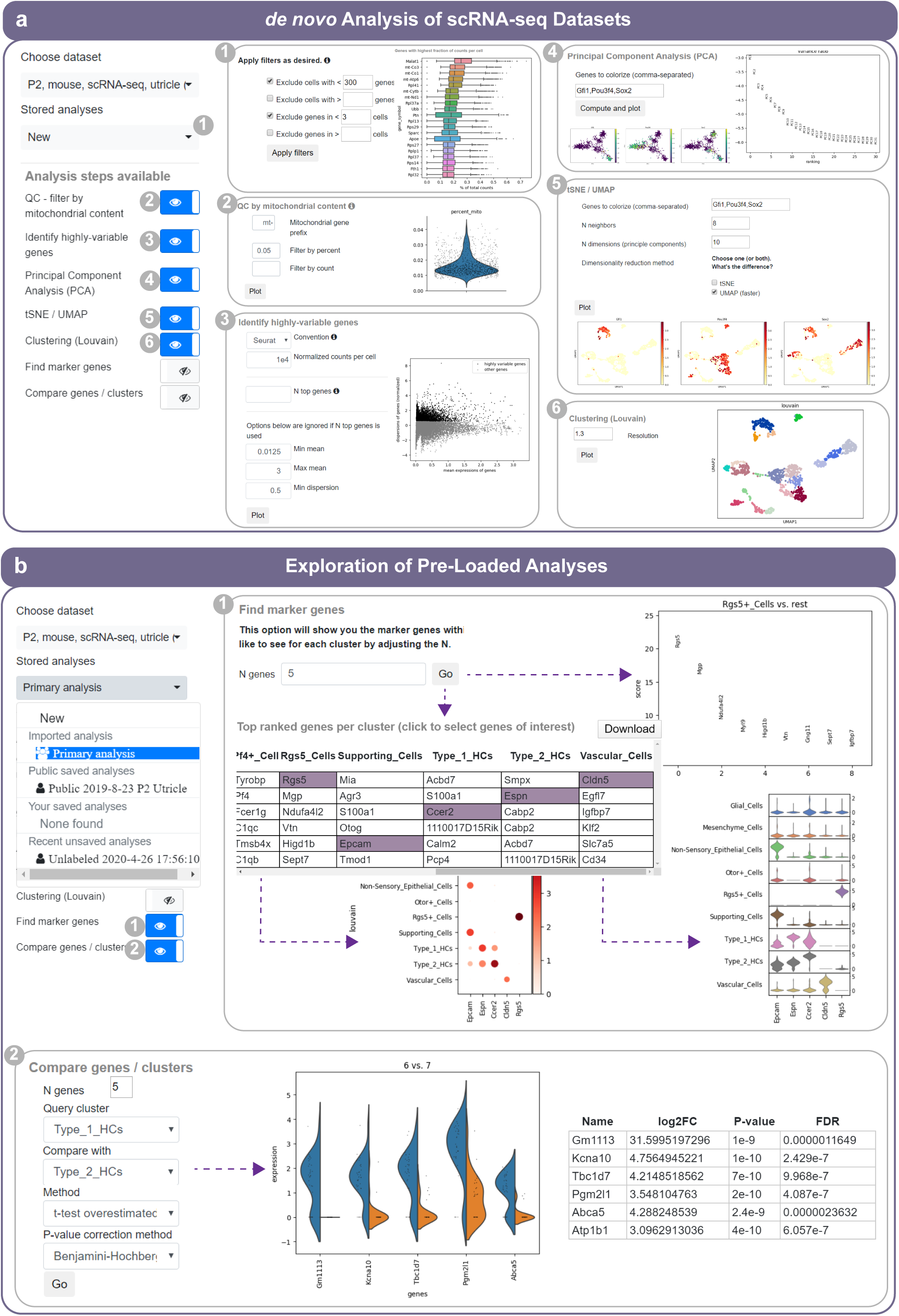
The gEAR scRNA-seq Workbench. **a**, *De novo* analysis of scRNA-seq data. The workbench, using toggle buttons, guides the users through the following steps: (1) Quality control including determining minimal and maximal genes per cell and based on the data structure and minimal number of cells with expression of a gene detected; (2) Ability to filter out cells with high mitochondrial gene content, which is common in dying cells; (3) Selection of highly variable genes for the dimensional reduction steps; (4) Principal component analysis (PCA) including graphic representation of the PC1 vs PC2 which can be colored by expression, and lists of the top genes contributing to chosen principal components; (5) Generating a tSNE/UMAP plots based on the results obtained in step 4; (6) Louvain clustering with renaming of the clusters after identifying marker genes for each cluster; as well as marker gene analysis and cluster comparison (shown in **b**). **b**, Working from a stored analysis allows the user to identify marker genes for the pre-defined clusters, and view their relative contributions for the corresponding cluster. Marker genes can be selected from the result table and their expression in every clusters can be plotted as violin and dot plots (1). Any two clusters can be compared directly using the compare clusters/genes toggle button (2).

### gEAR Genome Browser Integration

Transcriptional regulation of gene expression is reflected in changes to DNA accessibility and epigenetic marks, which are essential to deciphering gene regulatory networks. To visualize functional genomics data, we integrated gEAR with Epiviz^12^, allowing users to perform interactive visual and exploratory analysis of expression and epigenetic datasets within the same profile in gEAR. Users can upload epigenetic data to the gEAR portal and mark them as either public or private. After upload, users can customize the view of their data in the Epiviz-based tracks and perform integrative exploration of their epigenetic data aside datasets available on gEAR. An example Epiviz display is shown in **Fig. 5** for a dataset of reprogramming mouse embryonic fibroblasts (MEFs) to differentiate to cells similar to hair cells (the inner ear sensory cells)^19^. The expression of *Pou4f3* in the reprogrammed MEFs is shown in comparison to its expression in native hair cells and other cells types. In parallel, the DNA accessibility surrounding the *Pou4f3* locus is shown for native hair cells, MEFs and MEFs induced to differentiate to hair cells, allowing users to visually correlated changes in gene expression to DNA accessibility and immediately identify candidate regulatory elements. A classic workflow would include identifying marker genes or differentially expressed genes using the compare tool of the gEAR, generating a gene cart, and then exploring their genomic loci while verifying their differential expression all in the same page. Users could populate their page with additional dataset from their own laboratory or publicly available ones for further validation of the data. Finally, the Epiviz integration includes several methods to visualize and analyze signals and regions of interest, and users can also navigate from the integrated view to the standalone Epiviz viewer for additional features.

**Fig. 5:**
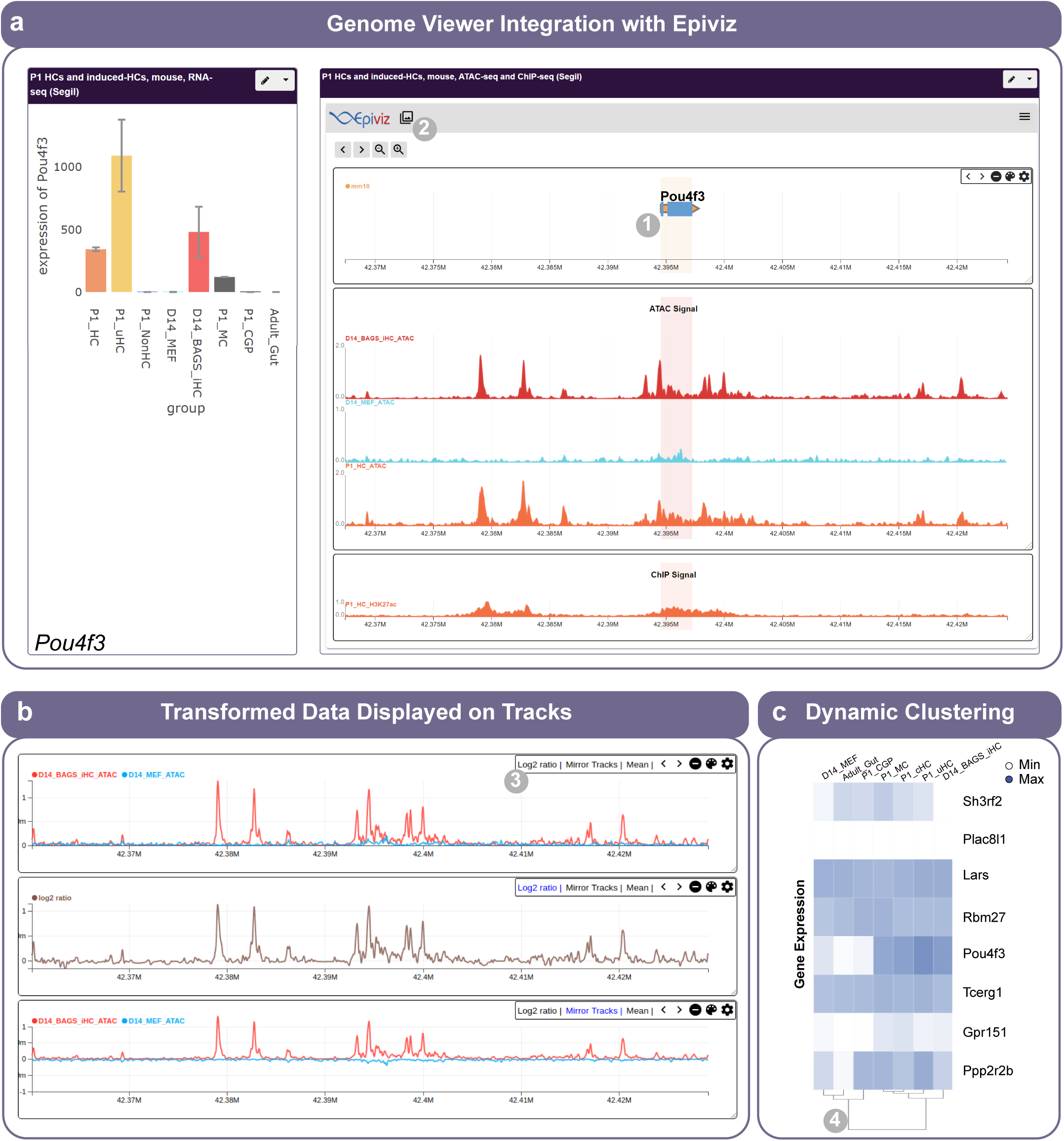
Integration of Epiviz in gEAR to visualize epigenetic data. **a**, An embedded view of a public dataset^19^ using the Epiviz browser within the gEAR portal. (from top to bottom) The first track displays the location of genes and their exons along the mm10 genome. The second track displays accessible regions inferred from ATAC-seq across samples. The last track shows H3k27ac ChIP-seq signal. Epiviz provides instant visual cues to the user when hovering over a track, highlighting this region across all the other tracks in the viewer (1). **b**, Epiviz provides multiple ways to visualize and transform data displayed in the track. For example (3) it provides options to apply log2 ratio, mirror tracks or calculate the average signal across all tracks in a visualization. In this case, the first track in this panel shows the raw ATAC signal for samples (induced hair cells (iHC) and mouse embryonic fibroblasts (MEF)), while the second track shows the log2 ratio of the signal between these two samples which identifies regions of differential chromatin accessibility. The third track mirrors the signal of the MEF sample to ease the visual comparison of the two samples. **c**, In the standalone mode, accessible via the pop-up icon (2), Epiviz also supports plot-based visualizations (Heatmaps and Scatter Plots) for interactive visualizations of gene expression data including dynamically rendered dendrograms as shown in (4).

### Dataset uploader

An important component of a system such as the gEAR is the ease with which users can upload complex data. The process of uploading data to the gEAR is interactive, guiding users through describing their data (and associated meta-data), choosing display preferences and selecting an access level (i.e., whether a dataset will be private or public). For RNA-seq data supported file formats include the Market Exchange (MEX) format, Microsoft Excel, and tab-separated text files. Templates are provided for each format as examples. Although some of these data files can be quite large, the validation and conversion to our internally used binary format finishes quickly. An example dataset with 19,591 genes and 5,253 cells was uploaded with the Dataset Uploader interface, validated and converted to H5AD format in 1.2 minutes. Similarly, the gEAR platform provides an interface to import epigenetic data by directly uploading the file or associating files from a trackhub configuration.

### Documentation, tutorial and helpdesk

It is critical that users have thorough documentation and in multiple forms. We have created extensive documentation in an online manual, guiding the user through the interface with screenshots and internal hyperlinks. We also provide training videos to demonstrate how to upload data and use the interface to exercise all features of the site. Finally, users can submit tickets via the helpdesk user interface, allowing for individual help where needed.

### Privacy, dataset sharing and data safety

The gEAR enables sharing of multi-omic data between collaborators. Users who upload datasets can post them publicly or store them for private use. Private datasets can be shared via “share URLs” e-mailed to other users. Users who receive a “share URL” can add those private datasets into their own profiles and view alongside their own datasets. There is a sharing manager so that the originator of the datasets can view and even revoke access to any given shared dataset, if needed.

### Open-source availability and user contributions

The entire portal code is written to be readily reused and themed for specific research interests. While the initial version of the gEAR was focused on the hearing research community, a clone of the entire site has also been created for brain researchers (nemoanalytics.org). Doing so requires only a checkout from GitHub, installation and then customization of a selection of style files to provide the desired theme. The gEAR code is available on GitHub (https://github.com/IGS/gEAR) and released under the GNU Affero General Public License v3.0. We have recently also added a plug-in framework so that users running their own portals can add additional, domain-specific functionality.

### Discovery using the gEAR portal – *Tmem178b* is an evolutionarily conserved hair cell expressed gene

The availability of numerous related datasets in the gEAR affords researchers in the auditory research community the ability to not only visualize gene expression or analyze one dataset at time, but also perform discovery research via rapid comparison between multiple related datasets within the gEAR. For example, here we chose to search for new evolutionarily conserved inner ear hair cell markers. We started our search with zebrafish using the workbench to identify hair cell makers in a homeostatic neuromast scRNA-seq dataset (**Fig. 6a**). We then chose three of the identified markers (*otofb, atoh1a* and *tmem178b*) to test for cell type-specificity of their orthologs (*Otof, Atoh1* and *Tmem178b*) using a mouse newborn cochlear epithelium scRNA-seq dataset, and found that all three genes were uniquely expressed in the cochlear hair cells (**Fig. 6b**). Of these, only *Tmem178b* was not previously described in the ear. Functional annotation indicates *Tmem178b* is a membrane bound protein, and a link-out to PubMed showed that little is known about this gene (**Fig. 6c**).Furthermore, the deafness gene annotation indicates that in humans *TMEM178B* is located on chromosome 7 (chr7:140,773,864-141,180,179) within the DFNB13 locus mapped in a family with autosomal recessive non-syndromic hearing loss^20^ (**Fig. 6c**).Further exploration showed that within the cochlea *Tmem178b* is uniquely expressed in hair cells from as early as embryonic day 14 (**Fig. 6d**). Viewing this gene within yet another dataset that recoded gene expression, now in sorted hair cells from the developing auditory and vestibular systems^21^, both validated this result and also revealed that while *Tmem178b* is specific to hair cells in the cochlea, it is expressed at even higher levels in the hair cells of the vestibular system (**Fig. 6e**). Inspection of other scRNA-seq datasets in the gEAR that include vestibular epithelia indicated that indeed within the vestibular system *Tmem178b* expression is unique to the hair cells with a higher expression in type II compared with type I hair cells (e.g., **Fig. 6f**). Moreover, this gene may play an important role in hair cell formation, as it is induced in regenerating hair cells as shown in another dataset where adult utricular supporting cells were transduced with *Atoh1* to generate hair cells (**Fig. 6g**). A discovery such as this without gEAR would require a researcher to download multiple data sets and analyze them using custom scripts which is not feasible for all biologists.

**Fig. 6:**
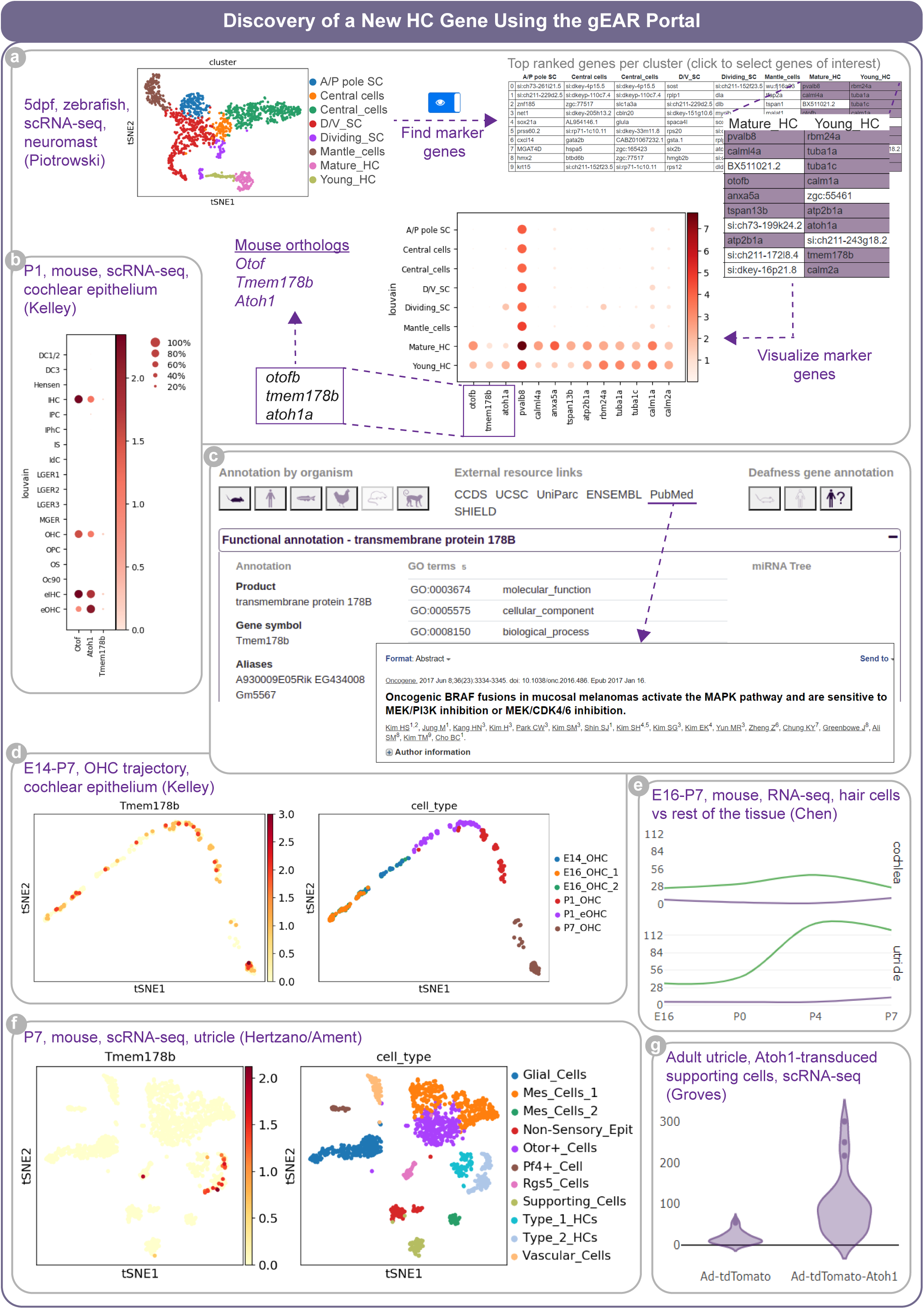
Discovery of a new HC gene using the gEAR portal. **a**, Identification of hair cell markers from a zebrafish scRNA-seq dataset^30^ using the scRNA-seq workbench. Three markers were selected, *atoh1a, otofb* and *tmem178b*. **b**, The mouse orthologs of the selected genes are also specific for HC when their expression is checked in a mouse scRNA-seq^5^ dataset. **c**, Functional annotation indicates that *Tmem178b* is a transmembrane protein and that the link to PubMed retrieves only one publication not related to hearing. The link to the deafness gene annotation reveals that the human *TMEM178B* is located on chromosome 7 within the DFNB13 locus^20^. **d**, A pre-computed trajectory analysis of a scRNA-seq dataset from cochlear epithelial cells of E14 to P7 mice^5^ reveals that *Tmem178b* is continuously expressed. **e**, Checking the expression of *Tmem178b* in a dataset featuring cochlear and vestibular HC^21^ indicates that the transcript is highly expressed in vestibular HC. **f**, Searching the expression in an utricle scRNA-seq dataset refines *Tmem178b* expression to vestibular Type II HC. **g**, Additionally, searching for *Tmem178b* in the “Regeneration” profile indicates that the gene is induced in supporting cells following ectopic expression of *Atoh1*^67^.

## DISCUSSION

While the abundance of functional genomics data is a blessing for many research fields, the size and complexity of most datasets make tools the biologist might traditionally use for analysis, no longer viable. Additionally, the rise of single cell multi-omics as a common discovery tool in biological sciences further challenges visualization, analysis and sharing of these data. Single cell-based multi-omic data are high-dimensional, require rigorous bioinformatics support, and even simply reading some large single-cell datasets into memory for processing far exceeds the hardware limitations of standard desktop-class machines.Therefore, the classically accepted approach for data sharing via deposition of raw data or even expression matrices into repositories such as the GEO^6^ provides only limited access to the data. With laboratories generating sometimes hundreds of datasets a year, and published data in most fields consisting of numerous high value multi-omic datasets – lack of meaningful access to these data is akin to requiring an expert to read manuscript text or an intermediate to visualize published figures.

This growing gap between high-value multi-omic data and the biologist as their natural consumer calls for new mechanisms for analysis, visualization, sharing and dissemination of data. To bridge this gap, a growing number of interactive analysis and visualization tools have been developed, ranging from online tools to R packages^11^. Common to all these tools, however, is their design to facilitate the visualization and/or analysis of one dataset/species at a time. Some of the tools require basic programming knowledge in Python (https://www.python.org/) or R (https://www.r-project.org/) and many require at least familiarity with the GitHub repository (https://github.com/) and the ability to install the tool on a local machine or a university managed server. While there is no doubt that to use data in a meaningful way one has to understand the concepts underlying their analysis, limiting the accessibility of today’s highest value discovery tools to a subgroup of biologists adept in bioinformatics depreciates their potential value.

Resource building is a complex task which requires careful consideration of users’ needs, skill sets, and the ecosystem of data they may be interested in. Not designed to replace bioinformatics expertise, the gEAR enables biologists and bioinformaticians alike to quickly add their data to a visual interface, compare results across experiments, and use analysis tools to quickly interrogate data. By giving users a shared space to add and view their data, providing mechanisms to share data privately with each other and adding the ability to take notes and comment on different datasets, communities are created.

We recognized that any analytical interface is limited by what the developers have created buttons for users to do, and that an increasing number of biologists are learning at least the basics of programming languages^22^. We therefore plan to allow users to further interrogate any datasets in a gEAR portal using integrated Jupyter notebook capability (https://jupyter-notebook.readthedocs.io/en/stable/). We also plan on continuous addition of new analysis tools, improvements in scalability via cloud node balancing, support for simultaneous multi-gene views and expansion of the workbench to support other data types. User surveys and feedback at conferences and workshops have guided development of the gEAR and will continue to drive future advancements.

The gEAR began with and has been embraced by the hearing research community. High-profile, hearing-related datasets have been made available in the gEAR even before publication, and links to gEAR as the primary method for accessing datasets are ubiquitous within talks/posters at professional conferences. Additionally, the gEAR has already been used for data dissemination^5,23–35^, hypothesis generation^36–44^ and validation of gene expression^29,30,45–57^ by numerous publications in the ear field. It has also been referenced by numerous groups as a useful tool^58–64^. Our team screens the literature on a regular basis, uploads and curates new datasets relevant to the hearing research community, and has thereby transformed the gEAR from a simple tool to a high value resource. The success of the portal as a resource is reflected in its over 900 users with active accounts, 400+ datasets and thousands of monthly visits for data interrogation, at the time this manuscript is written. The tools we have built in gEAR and our model of working together with the research community to identify and incorporate high-value datasets for visualization and analysis are readily extensible to many other fields.

---

**Supplementary Fig. 1:**
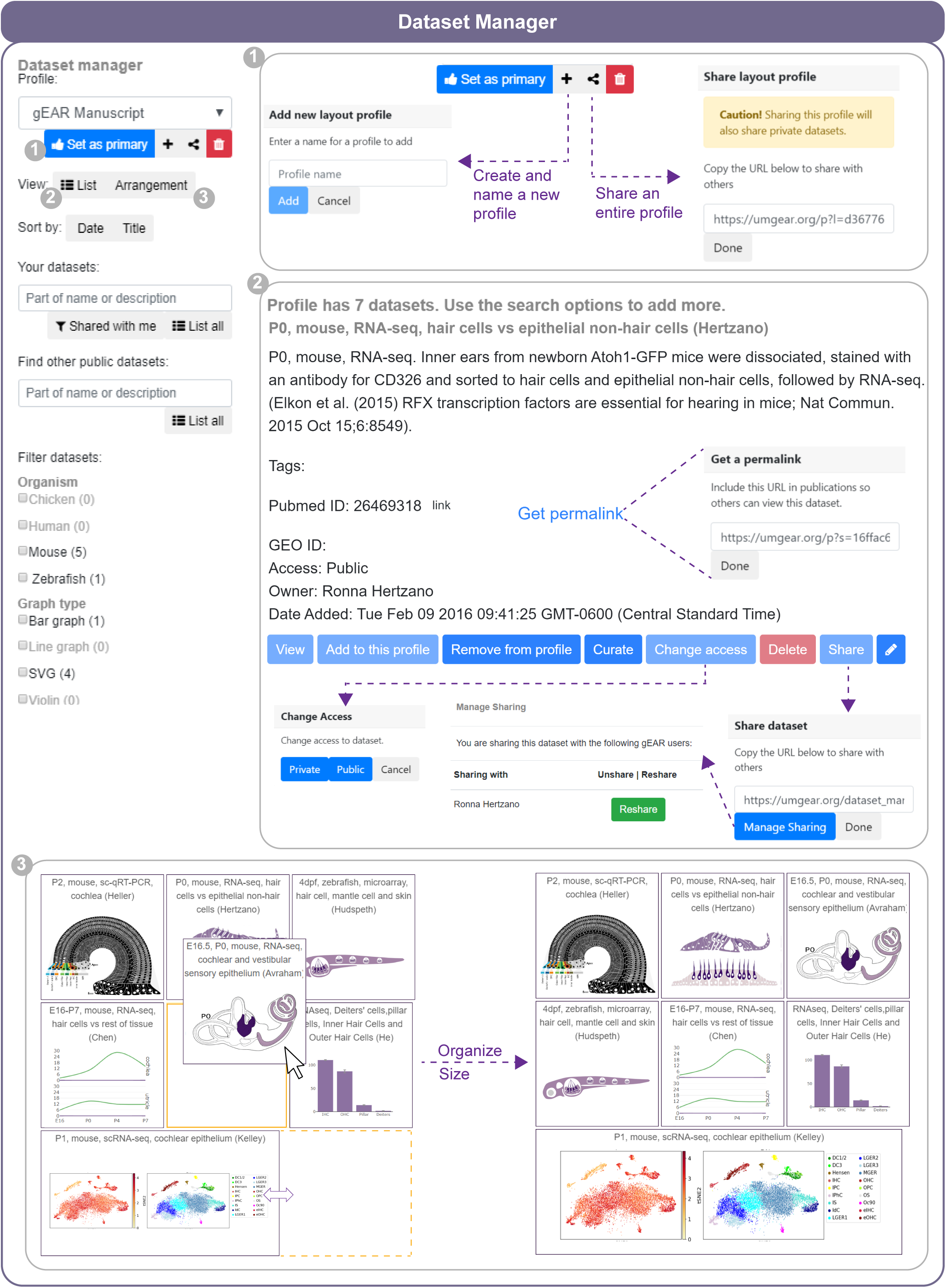
Dataset manager. The dataset manager allows users to identify datasets of interest and provides a set of filters fora more specific searches. (1) New profiles can be created and shared with collaborators. (2) The list view of the dataset manager contains information about the datasets and allows users to add/remove datasets from user-created profiles. A permalink can be created and added to publications and the datasets can be share with collaborators. The “Manage Sharing” button lists the people the dataset was shared with and allows datasets to be unshared. Lastly, the owner of a dataset can toggle the access between public and private. (3) The arrangement button allows users to configure a profile by changing the order or the size each dataset occupies on the grid for viewing.

**Supplementary Fig. 2:**
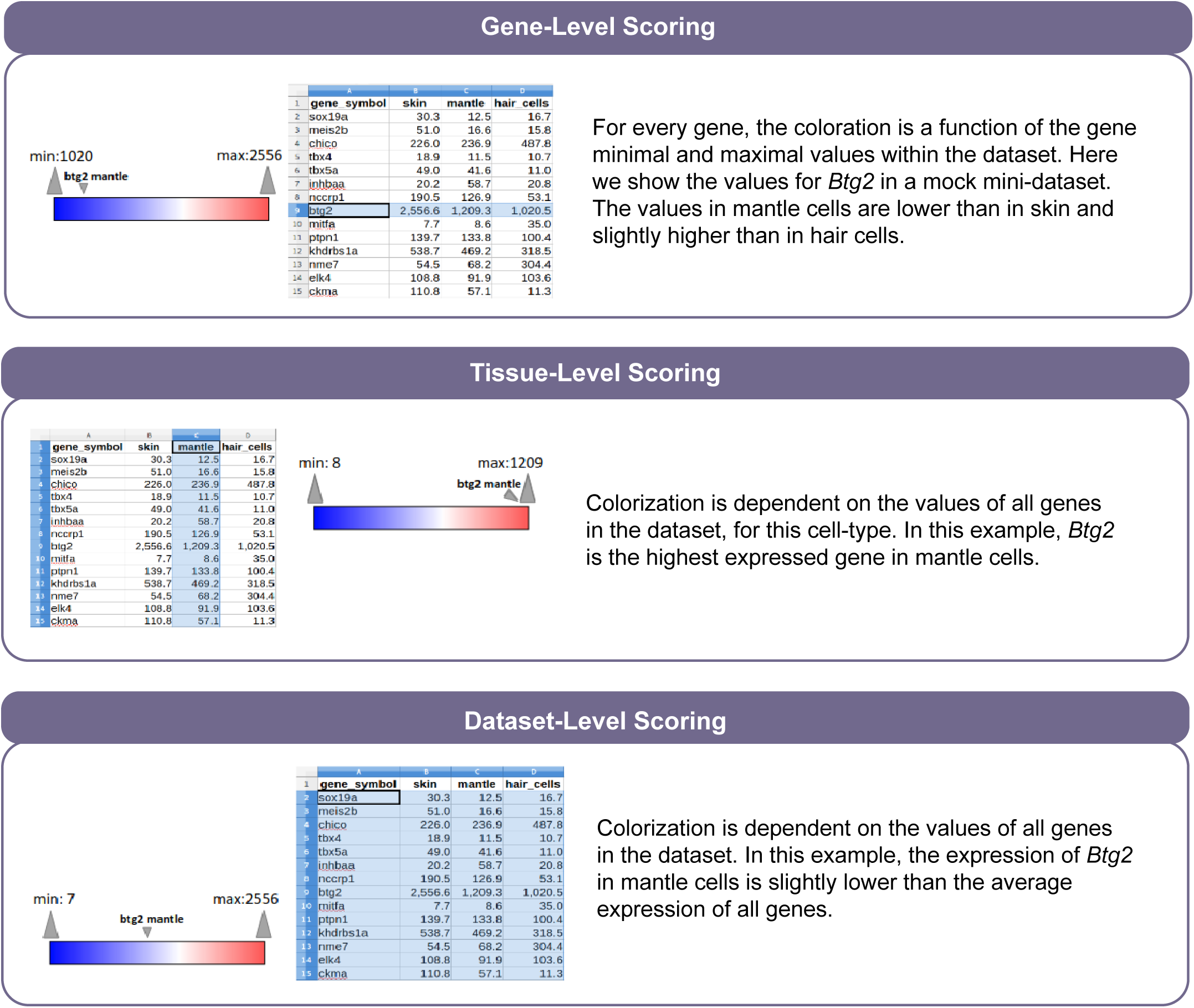
Color scoring for SVG figures. SVG figures have three types of coloring. Gene-level coloring (Top), Tissue-level coloring (Middle) and Dataset-level coloring (Bottom).

**Supplementary Fig. 3:**
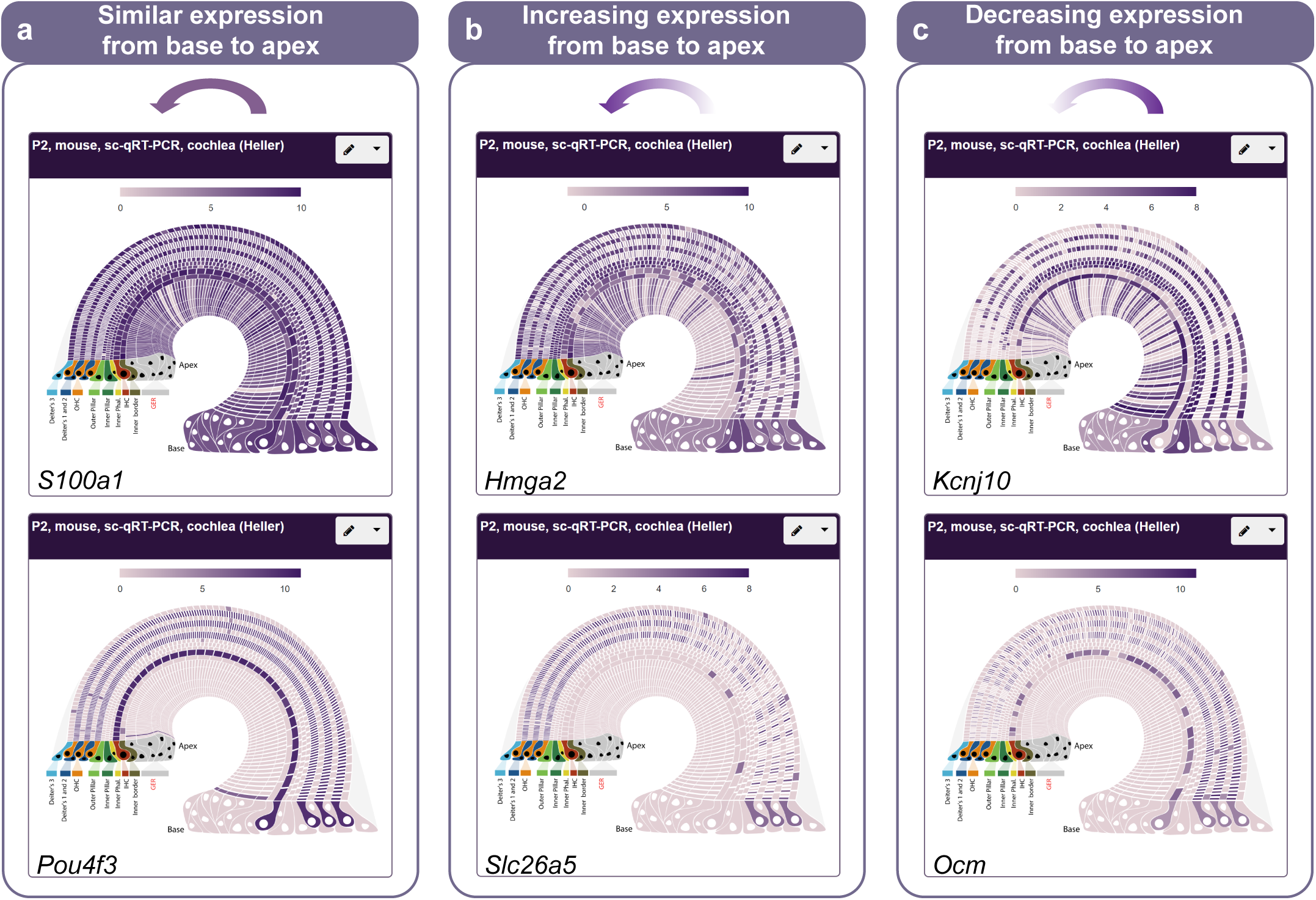
Scalable Vector Graphic (SVG) basal to apical representation of the organ of Corti. This is a graphical representation of an analysis of 877 single cells from an organ of Corti (hearing organ) from a 2-day old mouse. Gene expression of 192 genes was measured using RT-qPCR. Cell identity and basal to apical location along this organ of hearing were inferred from gene expression^68^. Here we generated an SVG for intuitive visualization of gene expression along the basal to apical axis as well as different cell types of the organ of Corti. **a**, *S100a1* (supporting cells) and *Pou4f3* (hair cells) have a constant expression from base to apex. **b**, *Hmga2* (supporting cells) and *Slc26a5* (hair cells) have a higher expression in the apex when compared to the base. **c**, *Kcnj10* (supporting cells) and *Ocm* (hair cells) show an increasing gradient of expression from apex to base.

**Supplementary Fig. 4:**
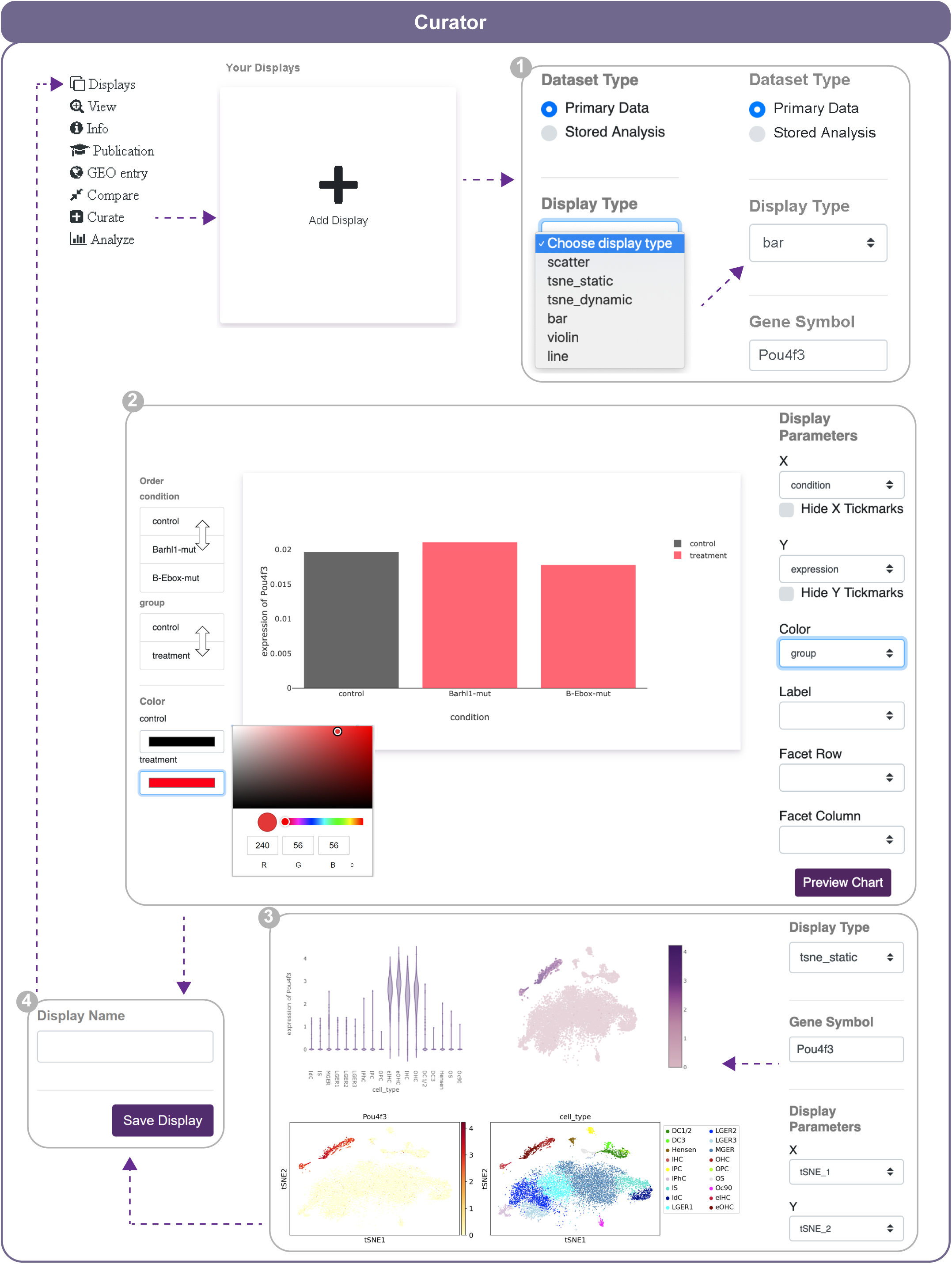
Display curator tool. The dataset curator allows users to create multiple display types for the same datasets. Different display options are listed depending on what the composition of the dataset enables (1). Choose a display type, enter a gene name, then choose display options (x-axis, y-axis, etc.) and an example plot will automatically update. More advanced options are available such as customized colors and gradients, column ordering and more (2 and 3). Once the created displays are saved (4), they can be retrieved from the dropdown menu associated with the corresponding dataset.

## ONLINE METHODS

### Dataset uploader for expression data

The gEAR was designed with a focus on a responsive user interface and flexibility in supporting different types of expression datasets. Registered users can upload their own datasets using the gEAR Uploader, an interactive web-based tool which serves as a guide through the upload process in a stepwise fashion. The upload process is the same for gene expression data (microarray, bulk RNA-seq and scRNA-seq). Input datasets begin as matrices in plain text grids or MS Excel spreadsheets. The most commonly uploaded variant of this format, and perhaps the richest and most straight-forward to generate, is the 3-tab input file set. The first file (expression.tab) is the expression matrix, with Ensembl IDs in the first column/key. Then there are metadata files for each row (genes.tab) and each subsequent column (observations.tab) of that matrix. The row metadata includes annotation attributes of each gene, such as gene symbol, while the columnar metadata can include information on cell type assignment, experimental condition, developmental stage, replicate information, or associated metrics such as standard deviation or P-value. Other common formats, such as the Market Exchange Format (MEX), are also accepted.

### Scalability

Once uploaded, the data are converted to a binary format called Hierarchical Data Format 5 (HDF5) [http://www.hdfgroup.org/HDF5/], following the schema defined by AnnData^18^. By using the indexed binary files on disk, rather than a relational database which was used in gEAR’s initial implementation, the system is now far more scalable. When the relational database was at its largest, querying expression data for 2 genes across 6 datasets took 7.5 minutes, which was reduced to 10.4 seconds after porting the schema to Google’s BigQuery. That wasn’t scalable to larger datasets, since BigQuery tables can’t support data matrices with more than 10,000 cells (). Finally, we ported to using binary HDF5 files (in H5AD format), both solving the scalability limitation and further reducing the same query speed to less than 2 seconds. Latency when searching a gene now depends only on the number of datasets present in the current profile (usually less than 12) rather than the number of datasets in the system altogether, since each dataset is an individual file and only those queried are even examined. This means a query will take the same few seconds whether the archive has 100 or 1000s of datasets uploaded.

### Epigenetic data

Epigenetic datasets can be stored within gEAR or hosted on any remote web server with sufficient features. While we currently accept user uploads of these large epigenetic datasets, we may require users to provide their own hosting in the future as resources allow. Epiviz uses an in-situ file-based query system (Epiviz File Server), that can directly query and transform indexed genomic files^69^. The server supports querying data by genomic location from both local and remotely hosted files. If the files are hosted remotely, Epiviz File Server uses HTTP range requests to only parse the minimum necessary bytes from a file to query data. This allows users to visualize epigenetic data hosted on their own servers without the need to host them on gEAR. In addition, Epiviz File Server also supports UCSC TrackHub configurations to integrate large datasets or repositories into the system. To visualize epigenetic data, we use the Epiviz component library^70^, an open source reusable and extensible data visualization library for functional genomic data. The embedded Epiviz browser supports a variety of charts to explore epigenetic data: block (linear and stacked) tracks for visualizing genomic regions of interest and line tracks (stacked, multi stacked) for visualizing signal (e.g., ATAC-seq, ChIP-seq and DNA methylation) data. This allows users on gEAR to choose an appropriate chart type to visualize their datasets. Hovering over a region in one visualization highlights this region in other tracks providing instant visual feedback to the user of the spatial overlap of different epigenetic data. These visualizations are developed using standards-based Web component framework, are highly customizable, reusable and can be integrated with most frameworks that support HTML^69,70^.

### Annotation and external links

All genes uploaded by users are linked to reference genes already stored within gEAR. These are primary annotations parsed from EMBL for *H. sapiens, M. musculus, D. rerio, G. gallus* and *R. norvegicus*, and updated regularly as EMBL issues new releases. We currently store all versioned releases for each organism back to Ensembl Release 83. Annotations displayed for each gene include gene product name, symbol, Enzyme Commission (EC) numbers and Gene Ontology (GO) terms. We have also mined available resources to provide direct link-outs for each gene to a large variety of external resources, including the UCSC (University of California Santa Cruz) genome browser^13^, SHIELD (Shared Harvard Inner-Ear Laboratory Database)^21^, CCDS (Consensus Coding Sequence) project^71^, Ensembl^72^, model organism databases, GeneCards^73^, Human Protein Atlas^74^, Pubmed, the DVD (Deafness Variation Database)^14^, the HHL (The Hereditary Hearing Loss Homepage)^75^, OMIM (Online Mendelian Inheritance in Men)^15^.

### Implementation details

The gEAR is written as a Python 3 application with client-based JavaScript libraries for creating responsive and interactive single page applications. The backend uses MySQL for storing annotation and user information, while the expression data and associated analyses are stored in individual binary HDF5 files. The API for accessing the databases uses Flask (https://palletsprojects.com/p/flask/) and the Scanpy^18^ module to process data, generating graphics using a variety of libraries including Matplotlib, Plotly and D3. The UI was created using a combination of standard HTML5, JavaScript and Vue.js. Each instance of gEAR is currently hosted on Google Cloud Platform nodes, allowing for quick resource allocation. There is also support for running a gEAR Portal within a Docker container.

## ACKNOWLEDGEMENTS

We are grateful to the members of the Hearing Restoration Project (HRP) for serving as the focus group in developing this tool and their continuous feedback. We thank Robert Marini for his technical assistance. This work was funded by the Hearing Restoration Project (R.H.); R01-DC013817 (R.H.); R24-MH114815 (R.H./O.R.W.); and R01-GM114267 (H.C.B).

## AUTHORS’ CONTRIBUTIONS

Portal design: J.O., B.G., A.M. and R.H. Data archive/infrastructure: J.O., B.G., J.K., M.C.K., H.C.B., S.A., C.C., B.H., O.W. and A.M. Graphic design: A.O.D. User interface: J.O., B.G., D.O. and A.M. User manual: J.O., Y.S., K.R., E.C. and R.H. DVD and Deafness gene/loci integration: H.A., J.O, and R.H. Providing unpublished data: K.R., M.C.K, M.W.K. and R.H. Project Management: R.L.C, A.M. and R.H. Data upload and analysis: J.O., D.O., Y.S., E.C., B.M., M.S.M. and R.H. Writing the manuscript: J.O., B.M., A.M. and R.H.

## REFERENCES

1. Shema, E., Bernstein, B. E. & Buenrostro, J. D. Single-cell epigenomics; review 2018. Nat. Genet. 51, 19–25 (2018).

2. Lonsdale, J. et al. The Genotype-Tissue Expression (GTEx) project. Nature Genetics 45, 580–585 (2013).

3. Yao, Z. et al. An integrated transcriptomic and epigenomic atlas of mouse primary motor cortex cell types. bioRxiv 2020.02.29.970558 (2020). doi: 10.1101/2020.02.29.970558

4. Kowalczyk, M. S. et al. Single-cell RNA-seq reveals changes in cell cycle and differentiation programs upon aging of hematopoietic stem cells. Genome Res. 25, 1860–1872 (2015).

5. Kolla, L. et al. Characterization of the development of the cochlear epithelium at the single cell level. Nat. Commun. 11, 2389 (2020).

6. Clough, E. & Barrett, T. The Gene Expression Omnibus database. in Methods in Molecular Biology 1418, 93–110 (Humana Press Inc., 2016).

7. National Center for Biotechnology Information. SRA Handbook. (2009).

8. Papatheodorou, I. et al. Expression Atlas update: from tissues to single cells. Nucleic Acids Res. 48, D77–D83 (2020).

9. Surujon, D. & Van Opijnen, T. ShinyOmics: Collaborative exploration of omics-data. BMC Bioinformatics 21, (2020).

10. Rohart, F., Gautier, B., Singh, A. & Lê Cao, K. A. mixOmics: An R package for ‘omics feature selection and multiple data integration. PLoS Comput. Biol. 13, (2017).

11. Çakır, B. et al. Comparison of visualisation tools for single-cell RNAseq data. bioRxiv doi: 10.1101/2020.01.24.918342

12. Chelaru, F., Smith, L., Goldstein, N. & Bravo, H. C. Epiviz: interactive visual analytics for functional genomics data. Nat. Methods 11, 938–40 (2014).

13. Haeussler, M. et al. The UCSC Genome Browser database: 2019 update. Nucleic Acids Res. 47, D853–D858 (2019).

14. Azaiez, H. et al. Genomic Landscape and Mutational Signatures of Deafness-Associated Genes. Am. J. Hum. Genet. 103, 484–497 (2018).

15. McKusick-Nathans Institute of Genetic Medicine, Johns Hopkins University (Baltimore, M. Online Mendelian Inheritance in Man, OMIM®. Available at: https://omim.org/.

16. Dickinson, M. E. et al. High-throughput discovery of novel developmental phenotypes. Nature 537, 508–514 (2016).

17. Satija, R., Farrell, J. A., Gennert, D., Schier, A. F. & Regev, A. Spatial reconstruction of single-cell gene expression data. Nat. Biotechnol. 33, 495–502 (2015).

18. Wolf, F. A., Angerer, P. & Theis, F. J. SCANPY: large-scale single-cell gene expression data analysis. Genome Biol. 19, 15 (2018).

19. Menendez, L. et al. Generation of inner ear hair cells by direct lineage conversion of primary somatic cells. Elife 9, 1–33 (2020).

20. Mustapha, M. et al. A sensorineural progressive autosomal recessive form of isolated deafness, DFNB13, maps to chromosome 7q34-q36. (1998).

21. Shen, J., Scheffer, D. I., Kwan, K. Y. & Corey, D. P. SHIELD: an integrative gene expression database for inner ear research. Database 2015, bav071 (2015).

22. Baker, M. Scientific computing: Code alert. Nature 541, 563–565 (2017).

23. Chessum, L. et al. Helios is a key transcriptional regulator of outer hair cell maturation. Nature 563, 696–700 (2018).

24. Hoa, M. et al. Characterizing Adult Cochlear Supporting Cell Transcriptional Diversity Using Single-Cell RNA-Seq: Validation in the Adult Mouse and Translational Implications for the Adult Human Cochlea. Front. Mol. Neurosci. 13, 13 (2020).

25. Malaiya, S. et al. Single-nucleus RNA-seq reveals dysregulation of striatal cell identity due to Huntington’s disease mutations. bioRxiv 2020.07.08.192880 (2020). doi: 10.1101/2020.07.08.192880

26. Engeln, M. et al. Individual differences in stereotypy and neuron subtype translatome with TrkB deletion. Mol. Psychiatry (2020). doi: 10.1038/s41380-020-0746-0

27. Sun, S. et al. Hair Cell Mechanotransduction Regulates Spontaneous Activity and Spiral Ganglion Subtype Specification in the Auditory System. Cell 174, 1247-1263.e15 (2018).

28. Matern, M. S. et al. Transcriptomic Profiling of Zebrafish Hair Cells Using RiboTag. Front. cell Dev. Biol. 6, 47 (2018).

29. Celaya, A. M. et al. Deficit of mitogen-activated protein kinase phosphatase 1 (DUSP1) accelerates progressive hearing loss. Elife 8, (2019).

30. Lush, M. E. et al. scRNA-Seq reveals distinct stem cell populations that drive hair cell regeneration after loss of Fgf and Notch signaling. Elife 8, e44431 (2019).

31. Korrapati, S. et al. Single Cell and Single Nucleus RNA-Seq Reveal Cellular Heterogeneity and Homeostatic Regulatory Networks in Adult Mouse Stria Vascularis. Front. Mol. Neurosci. 12, 316 (2019).

32. Ushakov, K. et al. Genome-wide identification and expression profiling of long non-coding RNAs in auditory and vestibular systems. Sci. Rep. 7, 8637 (2017).

33. Perl, K., Shamir, R. & Avraham, K. B. Computational analysis of mRNA expression profiling in the inner ear reveals candidate transcription factors associated with proliferation, differentiation, and deafness. Hum. Genomics 12, 30 (2018).

34. Burns, J. C., Kelly, M. C., Hoa, M., Morell, R. J. & Kelley, M. W. Single-cell RNA-Seq resolves cellular complexity in sensory organs from the neonatal inner ear. Nat. Commun. 6, 8557 (2015).

35. Wiwatpanit, T. et al. Trans-differentiation of outer hair cells into inner hair cells in the absence of INSM1. Nature 563, 691–695 (2018).

36. Majumder, P., Moore, P. A., Richardson, G. P. & Gale, J. E. Protecting Mammalian Hair Cells from Aminoglycoside-Toxicity: Assessing Phenoxybenzamine’s Potential. Front. Cell. Neurosci. 11, 94 (2017).

37. Requena, T., Gallego-Martinez, A. & Lopez-Escamez, J. A. Bioinformatic Integration of Molecular Networks and Major Pathways Involved in Mice Cochlear and Vestibular Supporting Cells. Front. Mol. Neurosci. 11, 108 (2018).

38. Barta, C. L. et al. RNA-seq transcriptomic analysis of adult zebrafish inner ear hair cells. Sci. data 5, 180005 (2018).

39. Li, Y. et al. Transcriptomes of cochlear inner and outer hair cells from adult mice. Sci. data 5, 180199 (2018).

40. Ingham, N. J. et al. Mouse screen reveals multiple new genes underlying mouse and human hearing loss. PLoS Biol. 17, e3000194 (2019).

41. Sadler, E. et al. Cell-Specific Transcriptional Responses to Heat Shock in the Mouse Utricle Epithelium. Front. Cell. Neurosci. 14, 123 (2020).

42. Ponniah, P. T. N. et al. Striatin is required for hearing and affects inner hair cells and ribbon synapses. bioRxiv 2020.03.11.987396 (2020). doi: 10.1101/2020.03.11.987396

43. Lewis, M. A., Domenico, F. Di, Ingham, N. J., Prosser, H. M. & Steel, K. P. Hearing impairment due to Mir183/96/182 mutations suggests both loss and gain of function effects. bioRxiv 579003 (2019). doi: 10.1101/579003

44. Jiang, F. et al. The ATPase mechanism of myosin 15, the molecular motor mutated in DFNB3 human deafness 1 2. bioRxiv 2020.06.17.155424 (2020). doi: 10.1101/2020.06.17.155424

45. Zheng, Q. Y. et al. An Age-Related Hearing Protection Locus on Chromosome 16 of BXD Strain Mice. Neural Plast. 2020, 8889264 (2020).

46. Ramzan, M. et al. Spectrum of genetic variants in moderate to severe sporadic hearing loss in Pakistan. Sci. Rep. 10, (2020).

47. Borse, V., Barton, M., Arndt, H., Kaur, T. & Warchol, M. E. Dynamic patterns of YAP1 expression and cellular localization in the developing and injured utricle. bioRxiv 2020.05.06.081323 (2020). doi: 10.1101/2020.05.06.081323

48. Friedman, T. B., Belyantseva, I. A. & Frolenkov, G. I. Myosins and Hearing. in Advances in experimental medicine and biology 1239, 317–330 (Adv Exp Med Biol, 2020).

49. Liu, H. et al. Transcription Co-Factor LBH Is Necessary for Maintenance of Stereocilia Bundles and Survival of Cochlear Hair Cells. bioRxiv 2020.05.13.093377 (2020). doi: 10.1101/2020.05.13.093377

50. Yousaf, R. et al. Mutations in Diphosphoinositol-Pentakisphosphate Kinase PPIP5K2 are associated with hearing loss in human and mouse. PLoS Genet. 14, e1007297 (2018).

51. Booth, K. T., Azaiez, H., Jahan, I., Smith, R. J. H. & Fritzsch, B. Intracellular Regulome Variability Along the Organ of Corti: Evidence, Approaches, Challenges, and Perspective. Front. Genet. 9, 156 (2018).

52. Lewis, M. A. et al. Whole exome sequencing in adult-onset hearing loss reveals a high load of predicted pathogenic variants in known deafness-associated genes and identifies new candidate genes. BMC Med. Genomics 11, 77 (2018).

53. Liu, H. et al. Cell-Specific Transcriptome Analysis Shows That Adult Pillar and Deiters’ Cells Express Genes Encoding Machinery for Specializations of Cochlear Hair Cells. Front. Mol. Neurosci. 11, 356 (2018).

54. Li, C. et al. Dysfunction of GRAP, encoding the GRB2-related adaptor protein, is linked to sensorineural hearing loss. Proc. Natl. Acad. Sci. U. S. A. 116, 1347–1352 (2019).

55. Bademci, G. et al. FOXF2 is required for cochlear development in humans and mice. Hum. Mol. Genet. 28, 1286–1297 (2019).

56. Jung, J. S. et al. Semaphorin-5B Controls Spiral Ganglion Neuron Branch Refinement during Development. J. Neurosci. 39, 6425–6438 (2019).

57. Dunbar, L. A. et al. Clarin-2 is essential for hearing by maintaining stereocilia integrity and function. EMBO Mol. Med. 11, e10288 (2019).

58. Farrell, B. & Bengtson, J. Scientist and data architect collaborate to curate and archive an inner ear electrophysiology data collection. PLoS One 14, (2019).

59. Hotchkiss et al. The Hearing Impairment Ontology: A Tool for Unifying Hearing Impairment Knowledge to Enhance Collaborative Research. Genes (Basel). 10, 960 (2019).

60. Hertzano, R., Gwilliam, K., Rose, K. P., Milon, B. & Matern, M. S. Cell Type-Specific Expression Analysis of the Inner Ear – A Technical Report. Laryngoscope Accepted, (2020).

61. Schrauwen, I. et al. Autosomal dominantly inherited greb1l variants in individuals with profound sensorineural hearing impairment. Genes (Basel). 11, 1–18 (2020).

62. Naz, S. & Friedman, T. B. Growth factor and receptor malfunctions associated with human genetic deafness. Clin. Genet. 97, 138–155 (2020).

63. Askew, C. & Chien, W. W. Adeno-associated virus gene replacement for recessive inner ear dysfunction: Progress and challenges. Hearing Research 107947 (2020). doi: 10.1016/j.heares.2020.107947

64. Pyle, M. P. & Hoa, M. Applications of single-cell sequencing for the field of otolaryngology: A contemporary review. Laryngoscope Investig. Otolaryngol. (2020). doi: 10.1002/lio2.388

65. Guo, Y., Zhang, P., Sheng, Q., Zhao, S. & Hackett, T. A. lncRNA expression in the auditory forebrain during postnatal development. Gene 593, 201–216 (2016).

66. Ranum, P. T. et al. Insights into the Biology of Hearing and Deafness Revealed by Single-Cell RNA Sequencing. Cell Rep. 26, 3160-3171.e3 (2019).

67. Jen, H. I. et al. Transcriptomic and epigenetic regulation of hair cell regeneration in the mouse utricle and its potentiation by Atoh1. Elife 8, (2019).

68. Waldhaus, J., Durruthy-Durruthy, R. & Heller, S. Quantitative High-Resolution Cellular Map of the Organ of Corti. Cell Rep. 11, 1385–1399 (2015).

69. Kancherla, J., Yang, Y., Chae, H. & Bravo, H. C. Epiviz File Server: Query, Transform and Interactively Explore Data from Indexed Genomic Files. Bioinformatics (2020). doi: 10.1093/bioinformatics/btaa591

70. Bravo, H. C., Kancherla, J., Zhang, A. & Gottfried, B. Epiviz Web Components: Reusable and extensible component library to visualize functional genomic datasets. F1000Research 7, (2018).

71. Pruitt, K. D. et al. The consensus coding sequence (CCDS) project: Identifying a common protein-coding gene set for the human and mouse genomes. Genome Res. 19, 1316–1323 (2009).

72. Cunningham, F. et al. Ensembl 2019. Nucleic Acids Res. 47, 745–751 (2019).

73. Stelzer, G. et al. The GeneCards suite: From gene data mining to disease genome sequence analyses. Curr. Protoc. Bioinforma. 2016, 1.30.1-1.30.33 (2016).

74. Uhlen, M. et al. Tissue-based map of the human proteome, Human Protein Atlas available from www.proteinatlas.org. Science (80-.). 347, 1260419–1260419 (2015).

75. Van Camp, G. & Smith, R. J. H. Hereditary Hearing Loss Homepage. Available at: https://hereditaryhearingloss.org.

